# *phylogenize:* correcting for phylogeny reveals genes associated with microbial distributions

**DOI:** 10.1101/425231

**Authors:** Patrick H. Bradley, Katherine S. Pollard

## Abstract

**Summary:** Phylogenetic comparative methods are powerful but presently under-utilized ways to identify microbial genes underlying differences in community composition. These methods help to identify functionally important genes because they test for associations beyond those expected when related microbes occupy similar environments. We present *phylogenize*, a pipeline with web, QIIME2, and R interfaces that allows researchers to perform phylogenetic regression on 16S amplicon and shotgun sequencing data and to visualize results. *phylogenize* applies broadly to both host-associated and environmental microbiomes. Using Human Microbiome Project and Earth Microbiome Project data, we show that *phylogenize* draws similar conclusions from 16S versus shotgun sequencing and reveals both known and candidate pathways associated with host colonization.

**Availability:** *phylogenize* is available at https://phylogenize.org and https://bitbucket.org/pbradz/phylogenize.

## Introduction

Shotgun and amplicon sequencing allow previously intractable microbial communities to be characterized and compared, but translating these comparisons into gene-level mechanisms remains difficult. Researchers typically correlate microbial gene abundance with environments using metagenomes, either from shotgun sequencing (Nayfach and Pollard, 2016) or imputed from amplicon sequences (Langille *et al*., 2013; Aßhauer *et al*., 2015). However, related microbes tend to both share genes and occupy similar environments, causing confounding. Phylogenetic methods can correct for such confounding in metagenomics data (Bradley *et al*., 2018), but are currently implemented only in command-line, computationally intensive software.

We developed *phylogenize*, a pipeline allowing researchers without specific expertise in phylogenetic regression to analyze their own data via the web, an R package (R Core Team, 2017), or the popular microbiome workflow tool QIIME2 (Bolyen *et al*., 2018). An important innovation specific to *phylogenize* is that input data can be shotgun metagenomes or 16S amplicon data, the latter being lower-cost and available for more environments. Using these taxonomic profiles and sample environments (i.e., sources), the tool returns genes associated with differences in community composition across environments.

## Overview

Users provide *phylogenize* with taxon abundances and sample annotations, in tabular or BIOM (McDonald *et al*., 2012) format. Shotgun data should be mapped to MIDAS species (Nayfach *et al*., 2016); amplicon data should be denoised to amplicon sequence variants (ASVs) with DADA2 or Deblur. *phylogenize* uses BURST (Al-Ghalith and Knights, 2017) to map ASVs to MIDAS species via individual PATRIC genomes (Wattam *et al*., 2014), using a default cutoff of 98.5% nucleotide identity (Rodriguez-R *et al*., 2018) and summing reads mapping to the same species. Taxa are linked to genes using MIDAS and PATRIC, and then gene presence is tested for association with one of two phenotypes: prevalence (frequency microbes are observed) or specificity (enrichment of microbes relative to other environments; see Bradley *et al*., 2018).

*phylogenize* is an R package with a QIIME2 wrapper written in Python and a web front-end written in Python with the Flask framework (Ronacher, 2018) and a Beanstalk-based queueing system (Rarick, 2014). *phylogenize* reports include interactive trees showing the phenotype’s phylogenetic distribution, heatmaps of significantly positively-associated genes, tables showing which SEED subsystems (Overbeek *et al*., 2005) are significantly enriched, and links to tab-delimited files containing complete results.

## Example Applications

### Human Microbiome Project

The Human Microbiome Project (HMP; Human Microbiome Project Consortium, 2012) collected both 16S amplicon and shotgun sequences from 16 body sites on 192 individuals. Shotgun data processing was previously described (Bradley *et al*., 2018). 6,577 amplicon samples were downloaded from the NCBI SRA and denoised with DADA2 (Callahan *et al*., 2016), combining reads from the same individual and site. We ran *phylogenize* on both data types to identify genes whose presence is associated with prevalence in the gut. Despite differing read depth and sequencing technology (454 versus Illumina), effect sizes for genes associated with gut prevalence were similar for amplicon and shotgun (0.339 ≤ *r* ≤ 0.601) and similar pathways were enriched (Figure 1A).

**Figure 1:**
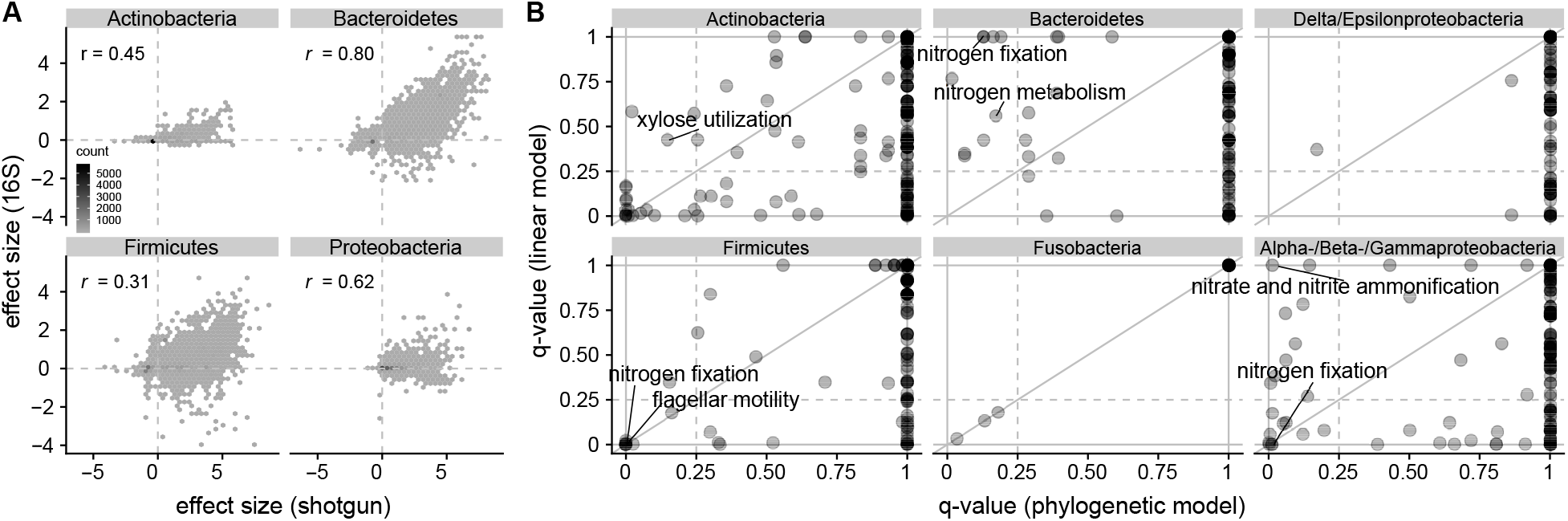
A. Effect sizes from HMP shotgun (x-axis) versus 16S amplicon (y-axis) data are correlated. Genes with *q* < 0.05 in one or both analyses shown with their Pearson correlation. Examples of SEED subsystems enriched for positively-associated genes with both data types include “Sporulation gene orphans” in Firmicutes (*q*_shotgun_ = 2.7 × 10^−22^, *q*_16S_ = 0.019) and “Type III, Type IV, Type VI, ESAT secretion systems” in Proteobacteria (*q*_shotgun_ = 1.69 × 10^−11^, *q*_16S_ = 2.23 × 10^−6^). B. SEED enrichments in EMP data using *phylogenize* (x-axis; 61 subsystems) or a linear model (y-axis; 202 subsystems). Many shared subsystems are relevant to a plant-associated lifestyle, such as nitrogen fixation (Mylona *et al*., 1995) and the metabolism of xylose (a pentose component of plant cell walls, Liu *et al*., 2015). Selected enrichments labeled; full list in Supplemental Table 1.

### Earth Microbiome Project

The Earth Microbiome Project (EMP) (Thompson *et al*., 2017) comprises 16S data from many biomes and habitats. Using the balanced subset of 2,000 samples processed using Deblur (Amir *et al*., 2017), we ran *phylogenize* and linear models (no phylogenetic correction) to identify genes whose presence is specific to plant rhizosphere compared to other environments. Linear models identified many more positively-associated genes (24,728 versus 7,490, *q* ≤ 0.05), but these discoveries were less enriched for processes known to be linked to plant rhizospheres (Figure 1B), suggesting dilution by false positives, as previously seen in HMP shotgun data and simulations (Bradley *et al*., 2018).

## Conclusion

Many microbes of interest to clinicians, ecologists, and microbiologists are poorly characterized or experimentally intractable. By making it easier to analyze either 16S or shotgun data with more precise statistical tools, *phylogenize* expands the toolkit for identifying mechanisms drivingdifferences in microbial community composition.

## Acknowledgements

Funding from the National Science Foundation [DMS-1069303 and DMS-1563159] and Gordon & Betty Moore Foundation [#3300].

